# Efference information flow during skill acquisition mediate its interaction with medical simulation technology

**DOI:** 10.1101/2021.09.19.460954

**Authors:** Anil Kamat, Basiel Makled, Jack Norfleet, Steven D. Schwaitzberg, Xavier Intes, Suvranu De, Anirban Dutta

## Abstract

Despite substantial progress towards establishing virtual reality (VR) simulators as a replacement for physical ones for skill training, its effect on the brain network during skill acquisition has not been well addressed. In this study, we employed portable optical neuroimaging technology and Granger causality approach to uncover the impact of the two medical simulation technologies on the directed functional brain network of the subjects with two different skill levels. The mobile brain-behavior relantionship was evaluated using functional near-infrared spectroscopy (fNIRS) while right-handed subjects performed well-established fundamentals of laparoscopic surgery (FLS) pattern cutting task. A multiple regression path analysis found that the cognitive-action information flow from the right prefrontal cortex to the supplementary motor area statistically significantly predicted the FLS task performance. Here, the skill level (expert vs novice) affected the cognitive-action information flow from the right prefrontal cortex and the efference copy information flow from the left primary motor cortex via supplementary motor area as hub to the cognitive-perception at the left prefrontal cortex, i.e., the action-preception link. The simulation technology (physical vs VR simulator) affected solely the cognitive-action information flow from the right prefrontal cortex to the left primary motor cortex; however, the interaction between the medical simulation technology) and the skill level affected the efference information flow from the left primary motor cortex to the right prefrontal cortex and from the supplementary motor area to the left prefrontal cortex. These discriminative findings are crucial since our VR simulator had face and construct validity. Therefore, our study highlighted the importance of efference information flow within the framework of the perception-action cycle when comparing medical simulation technology for visuomotor skill acquisition.

## Introduction

Virtual Reality (VR) technology is increasingly being used for motor skill training in medicine ^1^; however, the investigation of the perception-action cycle ^2^ in the VR compared to physical simulators is lacking. Specifically, there is a need to study the directional information flow that creates the sensorimotor mapping during motor skill acquisition. Here, the novice brain builds a multimodal cognitive-perceptual model ^3,4^ under perceptual load ^5^ to learn a goal-directed action during skill acquisition ^1^. Under the forward model concept ^6,7^, the central nervous system internally simulates the motor behavior in planning, control and learning such that the prediction of the sensory input is compared with the reafferent sensory input to determine discrepancy, e.g., at the frontal eye field for the visuosaccadic system ^8^. Christensen et al. ^9^ investigated the action-perception coupling based on the effect of action execution on action-perception that is postulated to be crucial for motor skill training ^1^. So, action for perception ^10^ that facilitates the closure of the perception-action cycle may be necessary during skill acquisition compared to learned skill performance by the experts. Then, the development of the sensorimotor mapping during skill acquisition is usually under variability “for all sets or series of observations that are non-constant and … non-stationary”^11^ including brain activation ^12^. Here, the inference about the state of the tool and the environment under noisy feedback will be made using a cognitive-perceptual that novices will develop for the closure of the perception-action cycle leading to a subjective sense of control ^13^. So, the understanding the variability in the brain and behavior in the structural and dynamic context of the perception-action cycle ^2^ can provide additional insights into skill acquisition. Here, the brain-behavior relationship in novices will differ from experts who have the cognitive-perceptual and may also have achieved certain motor skill “automaticity”^14^. To investigate the brain-behavior relationship in the context of the perception-action cycle ^2^, investigation of the directional information flow in the novice brain when compared to the expert brain may provide insights into the motor skill acquisition with various simulation technologies ^1^. For example, the dependence of perception on action ^10^ for motor skill acquisition may differ between the VR versus physical simulators. Such mobile brain-behavior investigation is now feasible due to the recent developments in portable brain imaging technologies ^15^, e.g., Nemani et al. ^16^ assessed bimanual motor skills using functional near-infrared spectroscopy (fNIRS) during laparoscopic surgery training.

Laparoscopic surgery training following the Fundamentals of Laparoscopic Surgery (FLS) is a common education and training module designed for medical residents, fellows, and the physician to provide them a set of basic surgical skills necessary to conduct laparoscopic surgery successfully. The FLS training is a joint education program between the Society of American Gastrointestinal Endoscopic Surgeons and the American College of Surgeon to establish box trainers (physical simulators) in standard surgical training curricula ^17^. FLS certification in general surgery in the USA uses five psychomotor tasks with increasing task complexity: (i) pegboard transfers, (ii) pattern cutting, (iii) placement of a ligating loop, (iv) suturing with extracorporeal knot tying, and (v) suturing with intracorporal knot tying. It was introduced to systemize training and evaluation of cognitive and psychomotor skills required to perform minimally invasive surgery. FLS is being used to measure and document those skills for medical practitioners where the understanding of the brain-behavior relationship is crucial for informed training and assessment ^18^, especially in the context of perception-action coupling ^19^ in physical versus VR simulators during laparoscopic surgery training ^20^. For example, surgeons rely on two-dimensional (2D) visualization of the three-dimensional (3D) surgical field at a reduced depth and tactile perception ^21^ where 3D vision has been shown to speed up laparoscopic training ^22^ possibly by reducing perceptual load ^5^ that is difficult to establish without the investigation of the information flow in the brain. Moreover, VR-driven sensorimotor stimulation may have harmful aftereffects ^23^ that need investigation of the brain-behavior relationship in the context of sensation weighting in the perception-action cycle ^19^. Furthermore, the understanding of the perception-action cycle in terms of differential sensation weighting can help in improving the design of the VR simulators in medicine in terms of not only visual and auditory but also kinesthetic and tactile feedback ^1,24^.

The FLS task performance (see Figure 1A) is graded based on the speed and accuracy related to psychomotor skills ^25^; however, not everyone can achieve proficiency ^26^. Here, we postulate that successful skill acquisition leads to an internal forward model ^27^ that can simulate the perceptual consequences of planned and executed motor commands. An intact action-perception coupling that is relevant for surgical skill acquisition has been shown to depend on the integrity of the cerebellum ^9^ that underpins the internal model ^28^. Then, the hierarchy of cognitive control during skill learning shows a rostrocaudal axis in the frontal lobe ^29^, where a shift from posterior to anterior is postulated to mediate progressively abstract, higher-order control expected in the experts. Here, the dorsolateral and ventrolateral prefrontal cortex (PFC) can be related to attention control, cognitive control, feature extraction, and formation of first-order relationships ^30,31,32,33^ relevant in novices. Specifically, dorsolateral PFC of the dorsal stream is more involved in the visual guidance of action while the ventrolateral PFC of the ventral stream is more involved in the recognition and conscious perception ^34^. Then, the supplementary motor area (SMA) and the premotor cortex are crucial for the coordination of bimanual movement ^35^ where SMA is crucial for complex spatiotemporal sequencing of movements ^36,37^ necessary in FLS tasks. Numerous functional magnetic resonance imaging (fMRI) and fNIRS studies have been published on skill learning ^21,38,39,40,41,42,43,44,45^; however, they have not systematically investigated the information flow ^46,47^ and its variability between experts and novices during surgical skill acquisition in physical versus VR simulators. Here, dynamic functional brain connectivity ^48^ can elucidate time-varying changes in brain activation and their dynamic reconfiguration ^49^. Specifically, the directed functional brain connectivity based on time-varying Granger causality analysis ^50^ can elucidate the directional information flow across brain regions in the context of perception-action coupling ^19^. Although fMRI studies have shown that the motor learning and transfer of learning from past experiences can be encoded by a large-scale brain network ^51,52^; however, fMRI is not suitable for mobile brain-behaviour studies. In this study, we used portable braim imaging with fNIRS that has limited spatial and depth sensitivity ^53^. Published fNIRS studies showed the involvement of the inferior parietal cortex, PFC, occipital cortex, and the sensorimotor areas, including premotor and primary motor cortex (PMC). In contrast, the fMRI studies showed additional activation of deeper brain structures, including the basal ganglia and cerebellum ^21^.

**Figure 1:**
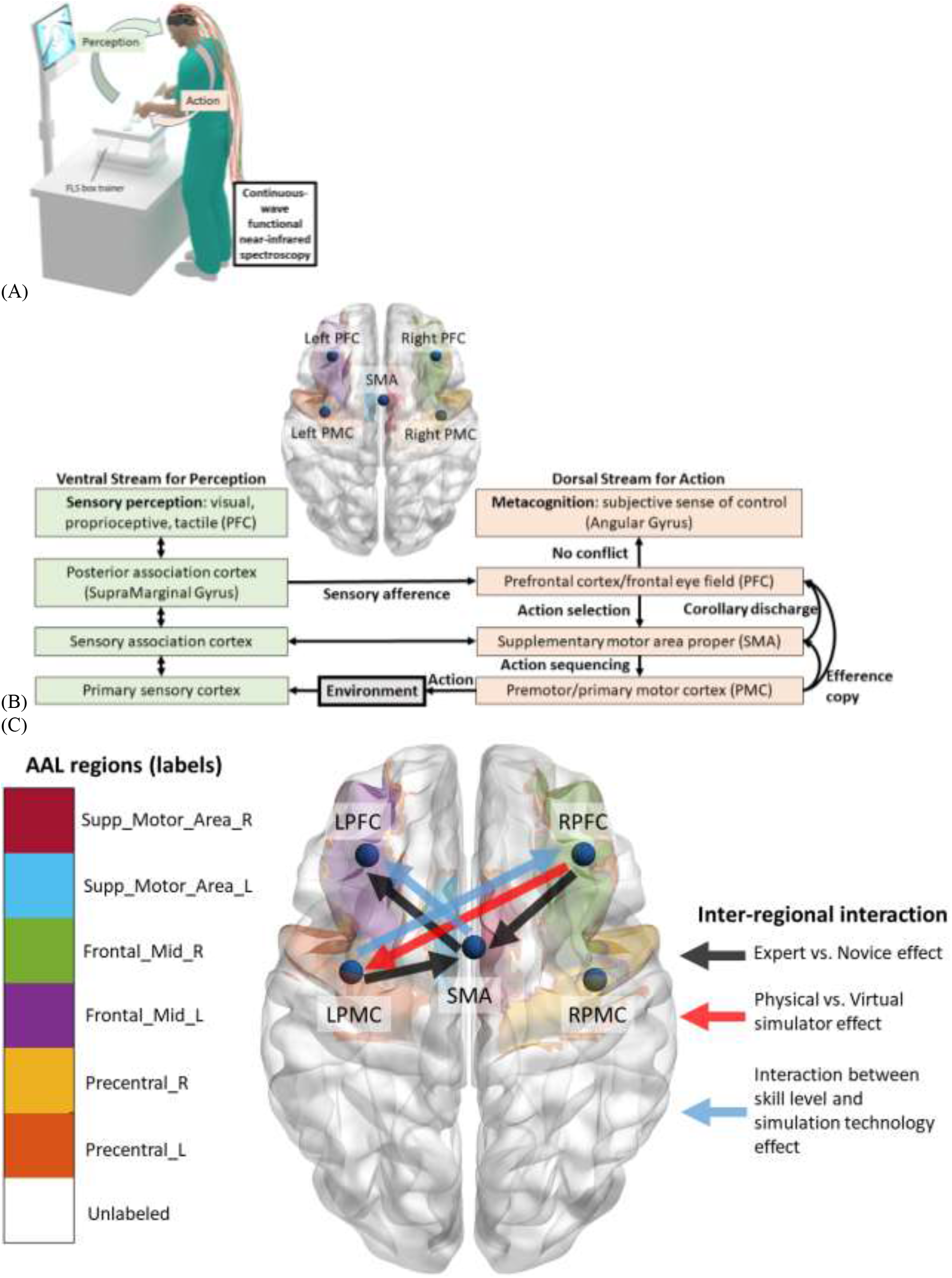
(A) Subject performing the bimanual FLS task while a continuous-wave spectrometer is used to simultaneously measure functional brain activation via functional near-infrared spectroscopy to capture the perception-action link to the surgical training. (B) Perception action link to surgical training in the physical and VR simulator environments. Our portable neuroimaging allowed investigation of the dorsal stream of action at the following brain regions, right PFC (RPFC), left PFC (LPFC), SMA, right PMC (RPMC), left PMC (LPMC) based on the sensitivity profile of our optode montage (Figure S1 in supplementary materials). (C) Automated anatomical labelling (AAL) of the brain regions (see Table S1), Supp_Motor_Area_R (SMA), Supp_Motor_Area_L (SMA), Frontal_Mod_R (RPFC), Frontal_Mod_L (LPFC), Precentral_R (RPMC), Precentral_L (LPMC) based on the optode sensitivity profile (Figure S1). Also, significant inter-regional directed functional brain connectivity are shown with colored arrows for the factors, the skill level (expert, novice), simulator technology (physical simulator, VR simulator), and their interaction.

In this study, we investigated the information flow ^54^ across the following brain regions, right PFC (RPFC), left PFC (LPFC), SMA, right PMC (RPMC), left PMC (LPMC) – see Figure 1B, based on the sensitivity profile of our optode montage (Figure S1 in supplementary materials). Figure 1B also shows the perception action link where our optode montage captured the dorsal stream for action starting from action selection in dorsolateral PFC to action sequencing in the SMA to action performance in the PMC. Then, the efference copy information from PMC is transmitted to the SMA and PFC while the corollary discharge from SMA is transmitted to the PFC. Here, we distinguished between efference copy versus collateral discharge based on whether motor action was transmitted versus motor plan for action-perception ^55^ in the PFC ^56^. Then, the action selection at the dorsolateral PFC and any conflict with the sensory reafference are monitored by the angular gyrus for a subjective sense of control ^13^ in the simulation environment. The ventral stream for perception of the sensory feedback from the environment at the primary sensory cortex flows to the sensory association cortex and then to the posterior association cortex (e.g. supramarginal gyrus) leading to conscious perception in the ventrolateral PFC. Here, PFC interacts through reciprocal and reentrant connections with different areas of posterior association cortex ^57^, including supramarginal gyrus, to integrate the information from sensory inputs and the actions ^58^ for action-perception ^55^. We found that the directed functional brain connectivity based on Granger causality (Figure S2 in supplementary materials) from the RPFC to SMA, LPMC to SMA, and SMA to LPFC during FLS task performance mediated the difference between experts and novices, as shown in Figure 1C alongwith the automated anatomical labelling (AAL) ^59^ of the brain regions with Montreal Neurological Institute (MNI) coordinates (see Table S1 in supplementary materials) based on the sensitivity profile (Figure S1 in supplementary materials). This presented SMA as the key junction ^60^ for the information flow that differentiated skill level (experts versus novices). Then, the difference between physical and VR simulators was captured by the directed functional brain connectivity from RPFC to LPMC mediated cognitive control that differentiated medical simulation technology (physical versus VR simulator). Also, an interaction between the medical simulation technology and the skill level was captured by the directed functional brain connectivity from LPMC to RPFC and SMA to LPFC (see Figure 1D) that can be related to efference copy and collateral discharge, respectively. Moreover, an interaction effect between the skill level and the simulator technology was found for the coefficient of variation (CoV) across trials of the directed functional brain connectivity from LPMC to RPMC. However, only the skill level and not the simulation technology was found to have a significant effect on the FLS task performance score and its CoV that highlighted the importance of portable brain imaging for capturing the perception-action cycle to evaluate medical simulation technology. Our prior work found wavelet coherence-based interhemispheric primary motor cortex connectivity and its CoV to be different between physical and VR simulators in the novices ^61^. In the current study, Granger causality and multiple regression approach identified directed information flow related to efference copy and corollary discharge linked to predictive internal signaling ^62^ within the framework of the perception-action cycle ^2^ that mediated the interaction between skill level and medical simulation technology.

## Results

### 1.1 Inter-regional directed functional brain connectivity

Repeated-measure two-way multivariate analysis of variance (two-way MANOVA) found a statistically significant effect of the skill level (expert, novice) on the inter-regional directed functional connectivity, RPFC to SMA (F(1,15)=6.045, p = .027; partial η2 = .287), LPMC to SMA (F(1,15)=7.892, p = .013; partial η2 = .345) and SMA to LPFC (F(1,15)=6.591, p = .021; partial η2 = .305). Also, two-way MANOVA found a statistically significant effect of the simulator technology (physical simulator, VR simulator) on the inter-regional directed functional connectivity, RPFC to LPMC (F(1,15)=6.002, p = .027; partial η2 = .286). Then, two-way MANOVA found a statistically significant effect of the interaction between the skill level and the simulator technology on the inter-regional directed functional connectivity, LPMC to RPFC (F(1,15)=8.523, p = .011; partial η2 = .362) and SMA to LPFC (F(1,15)=6.824, p = .02; partial η2 = .313). The details of the between-subject effects are presented in the Table S2 in supplementary materials.

Figure 2 shows the mean response for each factor (shown with colored arrows), adjusted for other variables in the model, i.e., the plot of estimated marginal means of the significant inter-regional directed functional brain connectivity. Figures 2(A), 2(B), 2(C) show the plot of estimated marginal means of the inter-regional directed functional brain connectivity affected by the skill level (expert, novice) where efference copy information flow from LPMC to SMA and the attentional control from RPFC to SMA is higher in expert than novices across both simulators. Then, Figure 2(D) shows the plot of estimated marginal means of the inter-regional directed functional brain connectivity affected by the simulator technology (physical simulator, VR simulator). Here, the higher inter-regional directed functional brain connectivity, RPFC to LPMC, in the VR simulator than physical simulator may be related to increased attentional processes ^63^ (or, attentional control) for fine motor control by the LPMC of the right-handed subjects since the RPFC optodes were over the right middle frontal gyrus (see Table S1 in supplementary materials). However, the inter-regional directed functional brain connectivity, RPFC to SMA, trended towards lower (Figure 2B) in the VR simulator than physical simulator that may underpin lesser attentional control of SMA. Also, the interregional directed functional connectivity, LPMC to SMA, trended towards higher (Figure 2A) in the VR simulator than physical simulator that may underpin stronger efference copy to SMA for modulating motor task. Then, the interregional directed functional connectivity, SMA to LPFC, considered the corollary discharge for action-perception ^55^ in PFC ^56^ trended towards lower in the VR simulator than the physical simulator for novice and attained a similar level as that of the expert – see Figure 2C. Here, VR simulator was novel for both the expert and the novice since experts were experienced with the physical simulator and human surgery. Therefore, an interaction between the skill level and the simulator technology was expected for the inter-regional directed functional connectivity, SMA to LPFC, as shown in Figure 2D. Moreover, the inter-regional directed functional connectivity, LPMC to RPFC, considered the efference copy for action-perception ^55^ in PFC ^56^, decreased in the VR simulator than the physical simulator for novice and attained a similar level as that of the expert – see Figure 2E. This may underpin similar level of internal forward model for expert and novice in the VR simulator.

**Figure 2:**
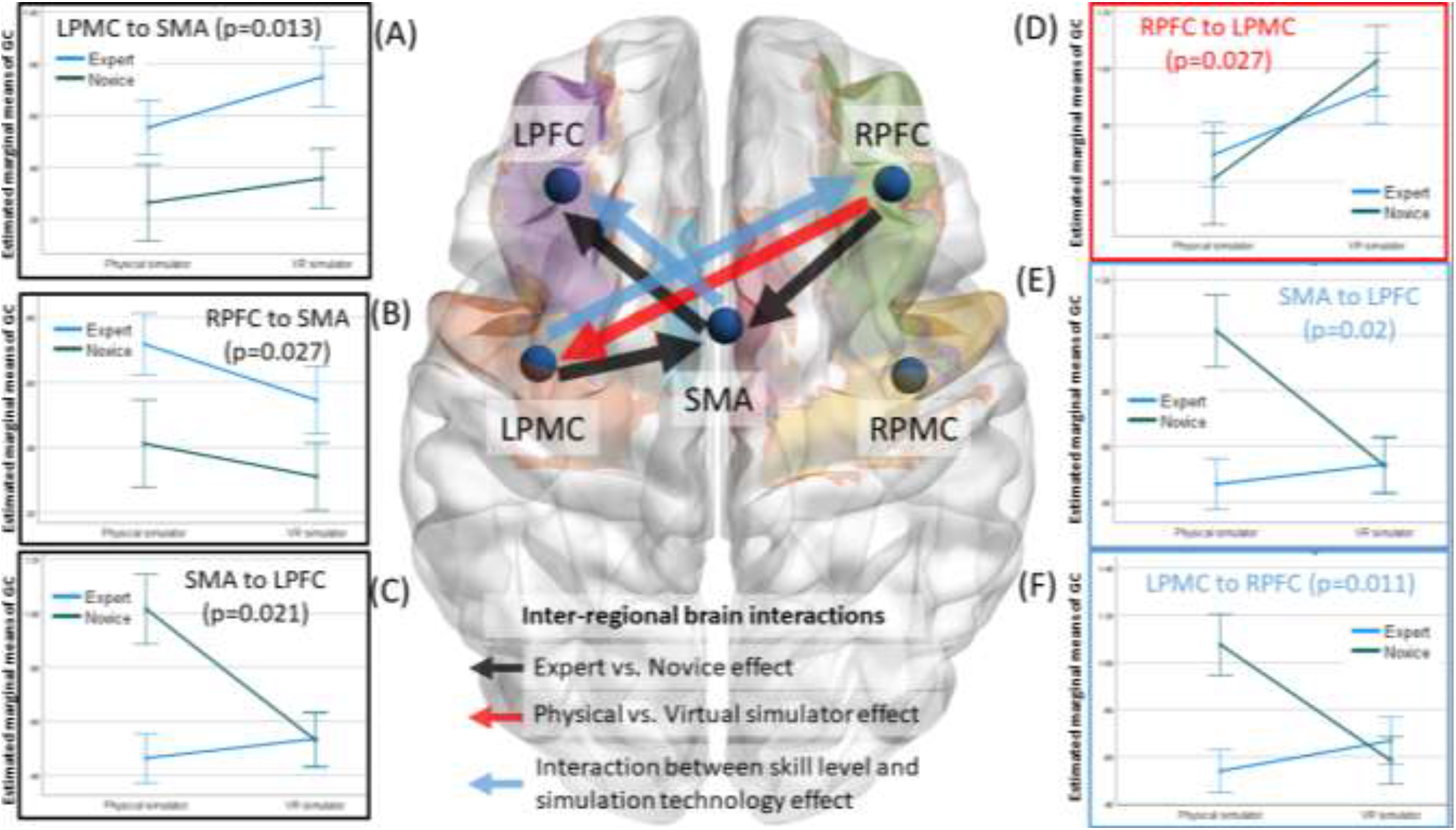
Plots of the estimated marginal means of the significant inter-regional directed functional brain connectivity. Error bars show standard error. (A) Significant (p=0.013) effect of the skill level (expert, novice) on LPMC to SMA directed functional connectivity. (B) Significant (p=0.027) effect of the skill level (expert, novice) on RPFC to SMA directed functional connectivity. (C) Significant (p=0.021) effect of the skill level (expert, novice) as well as (E) significant (p=0.02) effect interaction between skill level and the simulator technology on SMA to LPFC, directed functional connectivity. (D) Significant (p=0.027) effect of the simulator technology (physical simulator, VR simulator) on RPFC to LPMC directed functional connectivity. (F) Significant (p=0.011) effect of the interaction between skill level and the simulator technology on LPMC to RPFC directed functional connectivity.

### 2.2 FLS task performance score and coefficient of variation (CoV) of FLS task performance score

Repeated-measure two-way analysis of variance (ANOVA) found a statistically significant effect of the skill level on the FLS score (F(1,20)=12.786, p = .002; partial η2 = .39). Table S4 in supplementary materials presents the tests of between-subject effects. Two-way ANOVA also found a statistically significant effect of the skill level on the CoV of the FLS score (F(1,21)=4.370, p = .049; partial η2 = .172). Table S5 in supplementary materials presents the tests of between-subject effects. Figure 3A shows that the FLS score of the expert decreased in the VR simulator than the physical simulator since experts were experienced only with the physical simulator and human surgery. Then, Figure 3B shows that the CoV of the FLS score of experts increased in the VR simulator compared to the physical simulator.

**Figure 3:**
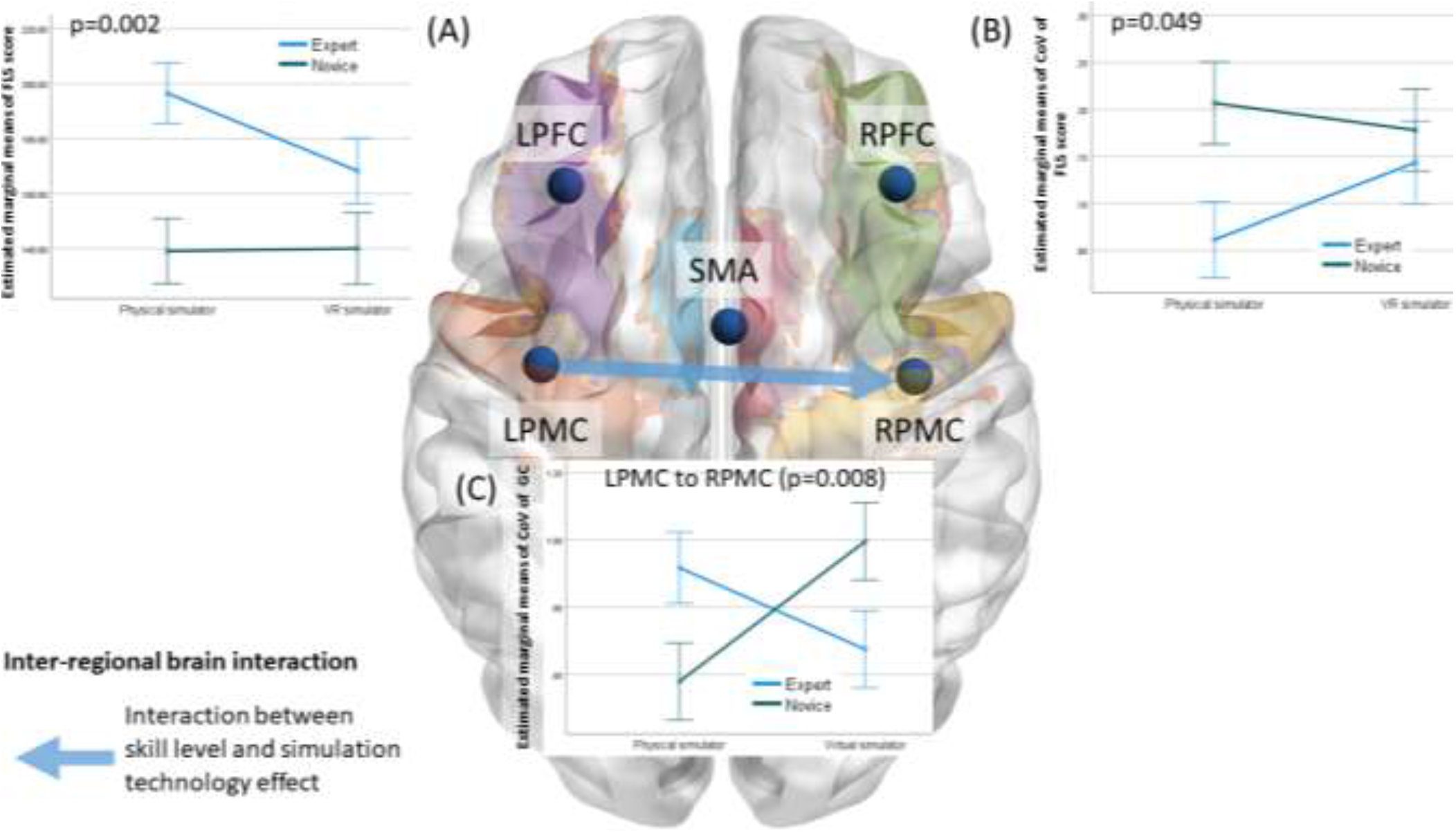
Estimated marginal means of (A) Significant (p=0.002) effect of skill level on FLS performance score and (B) Significant (p=0.049) effect of skill level on coefficient of variation (CoV) in FLS performance score. (C) Significant (p=0.008) effect of the interaction between skill level and the simulator technology on the coefficient of variation (CoV) of LPMC to RPMC directed functional connectivity. Error bars show standard error.

### 2.3 Coefficient of variation (CoV) of inter-regional directed functional brain connectivity

Two-way MANOVA found a statistically significant effect of the interaction between the skill level and the simulator technology on the CoV of the inter-regional directed functional connectivity, LPMC to RPMC (F(1,21)=8.561, p = .008; partial η2 = .29). Table S3 in supplementary materials presents the tests of between-subject effects. Figure 3C shows that the CoV of the inter-hemispheric directed functional connectivity, LPMC to RPMC, increased in the VR simulator than the physical simulator in novice while decreased in the expert. Here, CoV of the FLS score of experts increased in the VR simulator than the physical simulator (Figure 3B) so a decreased interhemispheric inhibition from LPMC to RPMC may underpin an increased CoV in performance in expert and vice versa in novice.

### 2.4 Brain-behavior relationships

Although the directed functional brain connectivity from the RPFC to SMA, LPMC to SMA, and SMA to LPFC mediated the difference between experts and novices while the difference between physical and VR simulators was captured by the directed functional brain connectivity from RPFC to LPMC (see Figure 2); however, the link between the directed functional brain connectivity and the FLS performance score can be elucidated based on multiple regression and path analysis (SPSS Amos, IBM, USA). Multiple regression (with backward elimination) analysis found that the FLS score was statistically significantly related to the inter-regional directed functional connectivity, RPFC to SMA, with F(2, 114) = 9, p < 0.001, R^2^ = .136 – see Figure 4A. Here, a significant partial regression (R^2^ = .136) of dependent variable, FLS score, with predictor, RPFC to SMA directed functional connectivity was found. Table S6 in supplementary materials presents the ANOVA results. Then, the regression weights from the path analysis (Figure S3 in supplementary materials) from factors (expert vs novice, physical vs VR simulator) to the directed functional brain connectivity (RPFC to LPMC, RPFC to SMA, LPMC to RPFC, LPMC to SMA, SMA to LPFC) to the FLS performance (FLS score) are shown in Table 1.

**Figure 4:**
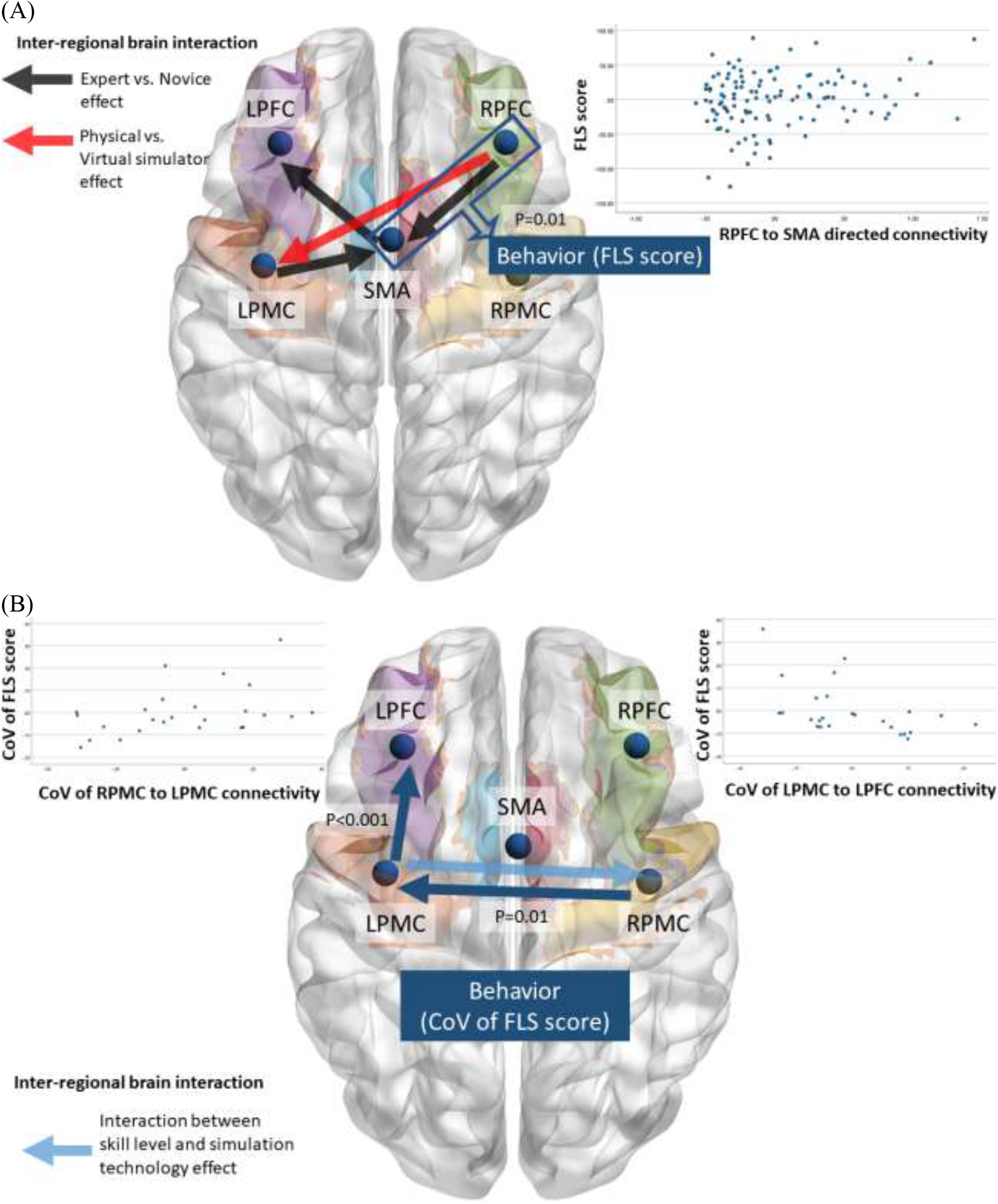
Brain-behavior relationships for FLS score and the coefficient of variation (CoV) of FLS score. (A) Interregional directed functional brain connectivity, RPFC to SMA, as a significant (p=0.01) predictor of the FLS score where RPFC to SMA functional connectivity is significantly (p<0.05) affected by the skill level (Expert vs Novice). The plot shows the partial regression of the inter-regional directed functional brain connectivity, RPFC to SMA, as a predictor of the FLS score. (B) CoV of the inter-regional directed functional brain connectivity, RPMC to LPMC (p=0.01) and LPMC to LPFC (p<0.001), as significant predictors for the CoV of the FLS score. CoV of the LPMC to RPMC functional connectivity is significantly (p<0.05) affected by the interation between the skill level and the simulation technology. The plots show the partial regression of the CoV of inter-regional directed functional brain connectivity, RPMC to LPMC and LPMC to LPFC, as a predictor of the CoV of the FLS score.

**Table 1:** Regression weight estimates for path analysis of FLS score (p<0.05 significant, grayed). Standard Errors (S.E.) and Critical Ratios (C.R.) are also shown besides p-value (P)

| Dependent var. |  | Independent var. | Weight Estimate | S.E. | C.R. | P |
| --- | --- | --- | --- | --- | --- | --- |
| RPFC to LPMC | <--- | expert vs novice | 0.02 | 0.12 | 0.13 | 0.89 |
| RPFC to SMA | <--- | expert vs novice | -0.23 | 0.08 | -2.94 | 0.00 |
| LPMC to RPFC | <--- | expert vs novice | 0.15 | 0.11 | 1.47 | 0.14 |
| LPMC to SMA | <--- | expert vs novice | -0.21 | 0.09 | -2.45 | 0.01 |
| SMA to LPFC | <--- | expert vs novice | 0.22 | 0.09 | 2.32 | 0.02 |
| RPFC to LPMC | <--- | physical vs VR simulator | 0.23 | 0.12 | 1.93 | 0.05 |
| RPFC to SMA | <--- | physical vs VR simulator | -0.13 | 0.08 | -1.63 | 0.10 |
| LPMC to RPFC | <--- | physical vs VR simulator | -0.09 | 0.11 | -0.84 | 0.40 |
| LPMC to SMA | <--- | physical vs VR simulator | 0.06 | 0.09 | 0.64 | 0.52 |
| SMA to LPFC | <--- | physical vs VR simulator | -0.12 | 0.09 | -1.26 | 0.21 |
| FLS score | <--- | RPFC to LPMC | -7.11 | 5.83 | -1.22 | 0.22 |
| FLS score | <--- | RPFC to SMA | 21.01 | 8.43 | 2.49 | 0.01 |
| FLS score | <--- | LPMC to RPFC | 3.66 | 6.50 | 0.56 | 0.57 |
| FLS score | <--- | LPMC to SMA | 3.13 | 7.89 | 0.40 | 0.69 |
| FLS score | <--- | SMA to LPFC | -1.16 | 7.15 | -0.16 | 0.87 |

When investigating brain-behavior relationships in terms of CoV, multiple regression (with backward elimination) analysis found that the CoV of the FLS score was statistically significantly related to the CoV of the inter-regional directed functional connectivity, RPMC to LPMC, LPMC to LPFC, with F(2, 22) = 3.912, p = 0.035, R^2^ = .262 – see Figure 4B. Table S7 in supplementary materials presents the ANOVA results. Table S7 in supplementary materials presents the ANOVA results. Then, the regression weights from the path analysis (Figure S4 in supplementary materials) from factors (expert vs novice, physical vs VR simulator) to the CoV of the directed functional brain connectivity (RPFC to LPMC, RPFC to SMA, LPMC to RPFC, LPMC to SMA, SMA to LPFC) to the CoV of the FLS performance (FLS CoV) are shown in Table 2. Here, the regression weight estimates for CoV of the RPMC to LPMC was 0.21 and for the CoV of the LPMC to LPFC was −0.30, where they both were statistically significant, p < .05.

**Table 2:**
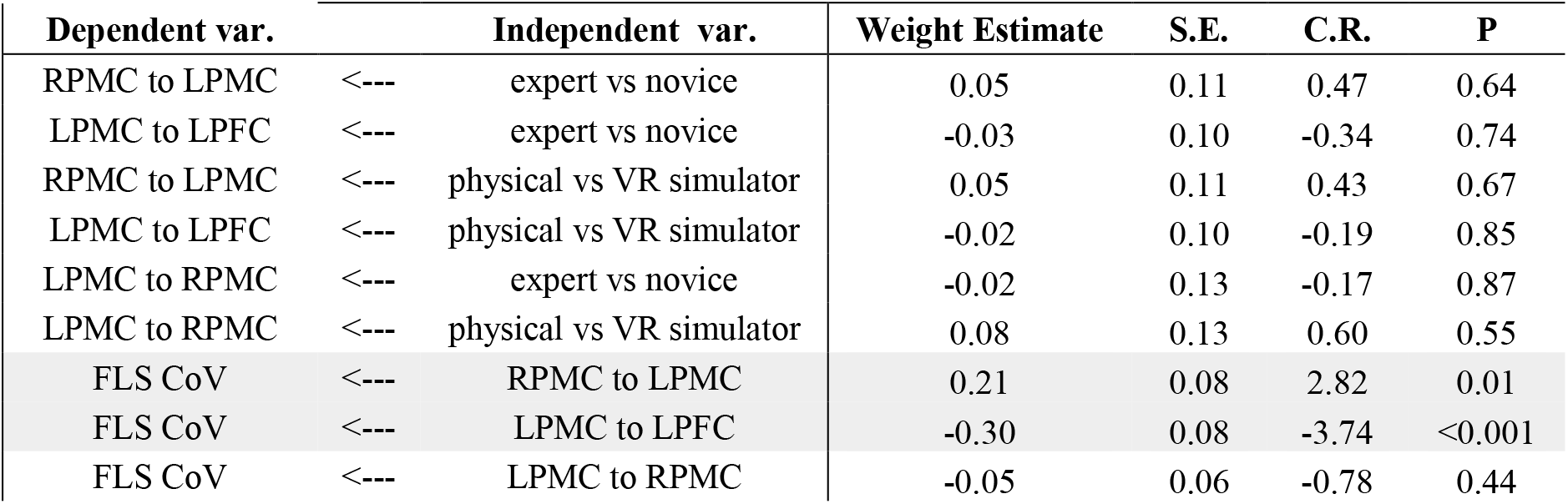
Regression weight estimates for path analysis of CoV of the FLS score (p<0.05 significant, grayed). Standard Errors (S.E.) and Critical Ratios (C.R.) are also shown besides p-value (P)

## Discussion

Our prior work found coupling between the PFC and the SMA using wavelet coherence-based functional connectivity metric that distinguished experts from novices during FLS pattern cutting in a physical simulator ^64^; however, the directionality of the information flow was not investigated. Since the brain-behavior relationship in terms of the directionality of the functional brain connectivity and its CoV was not investigated across physical and VR simulators; therefore, we applied spectral Granger causality ^50^ to determine the directional information flow in the brain networks and its CoV in physical and VR simulators in this study. Specifically, the SMA region was considered a key structure ^60^ for directed information flow from the LPFC, RPFC, LPMC, RPMC brain regions during the bimanual sequence operations task ^65,37,66,67,^ as shown in Figure 1, illustrating the perception-action link to FLS surgical training. SMA is a crucial region for interlimb coordination as well as eye-hand coordination ^68,69,70,71^ that is critical for perception-action coupling of the temporal organization and bimanual movement execution ^65,37,66,67^. So, the top-down executive control of the SMA is expected to differ ^14^ between experts and novices, where PFC ^72^ is known to have higher relevance in novices in facilitating training-induced task performance ^16^. Therefore, we applied a directed functional connectivity approach ^48^ to the fNIRS time series to capture the cascading directional processing of the goal-directed action ^73^, as shown by the dorsal stream of action in the Figure 1B. Here, sliding-window Granger causality provided a tool for identifying directed functional interactions from the fNIRS time series data that did not assume a static functional brain network across the whole FLS task, so also captured the CoV across repeated trials of the FLS task.

In this study, we found that portable brain imaging for brain-behavior modeling can evaluate medical simulation technology in terms of its interaction with the skill level within the context of the perception-action cycle ^2^. Specifically, prior work ^74^ had established the face and construct validity of the VR simulator used in this study; however, our investigation of the directed information flow of the brain regions relevant in perception-action coupling revealed the difference between physical and VR simulators. This difference was captured by the directed functional brain connectivity from RPFC to LPMC during FLS task performance that was higher in VR simulator than physical simulator for both expert and novice where PFC subserves cognitive control ^75^ and attentional processes ^63^ that are necessary for fine motor control. However, the distinguishing directed information flow for skill level as a predictor of FLS performance was found to be from RPFC to SMA (see Figure 4A) that trended towards lower in VR simulator than physical simulator (see Figure 2B). Jenkins et al. ^76^ have shown that PFC activation is associated with the learning of new sequence tasks while the lateral premotor cortex is more activated during new learning and the SMA is more activated during the performance of pre-learned sequence. Therefore, descent of the information flow from PFC to premotor/motor cortex ^77,78^ is expected in VR simulator that was novel for both the expert and the novice. Then, an interaction between the medical simulation technology (physical vs VR simulator) and the skill level (experts vs novices) was captured by the directed functional brain connectivity from LPMC to RPFC and SMA to LPFC (see Figure 2F and 2E) that are related to efference copy and corollary discharge information flow respectively (see Figure 1B). Here, SMA is known to contribute to the prediction of the sensory consequences of movement ^79^, which is expected when the internal forward model is available (e.g., in experts in the physical simulator). Therefore, corollary discharge ^80^ from SMA to the PFC is expected in the experts who have experienced physical simulators and human surgery for the cognitive control of bimanual movement ^36,37^. However, the VR simulator was novel for both the experts and the novices so the corollary discharge ^80^ from SMA to the PFC was found to drop from physical to VR simulator in the experts and was found comparable to the novices in the VR simulator (see Figure 2E). Furthermore, the efference copy from the LPMC to RPFC is postulated to be related to the functional coupling of the prefrontal and the premotor/motor areas that are expected during cognitive manipulation ^81^. Here, an increased cognitive manipulation ^81^ is postulated in the VR simulator when compared to the physical simulator due to the changes in the sensory signals ^55^ for both the experts and the novices (both inexperienced in VR), i.e., an increased information flow from RPFC to LPMC in the VR simulator (see Figure 2D). Also, the efference copy from the LPMC to RPFC was found to drop from physical to VR simulator in the experts due to a lack of internal model and was found comparable to the novices in VR (see Figure 2F).

Our study established a brain and behavior relationship based on fNIRS data that provided a portable, low-cost brain-imaging tool to compare task-related brain activation and functional brain networks in ambulant subjects for validating medical simulation technology for laparoscopic surgical tasks. Such validation based on the brain and behavior relationship is crucial since psychomotor skill learning or adjusting to changes in the environment, e.g., physical versus VR environment, requires the activation of brain regions and brain networks related to the perception-action cycle ^82,73^. Here (see Figure 3A and 3B), only the skill level and not the simulator technology was found to have a significant effect on the FLS score and its CoV (across trials), which aligns with the prior work ^74^ that established the face and construct validity of the VR simulator. Specifically, the expert had higher FLS score (Figure 3A) and lower CoV (Figure 3B) than novice in the physical simulator; however, in the VR simulator the expert without VR experience trended towards similar level as the novice. Motor skill learning requires sensory processing and transduction into a series of goal-directed actions where motor variability influencing task performance can shape motor learning ^83,84^ and motor variability typically tends to decrease with practice ^85^. So, subjects are expected to learn to avoid the influence of motor variability on goal-directed task performance ^83,84^ as observed in the experts with smaller CoV in the task performance (FLS score) in the physical simulator than novice while having similar CoV in the inexperienced VR simulator (Figure 3B). Then, the variability (CoV) in the task performance (FLS score) was significantly related to the variability (CoV) in the directed functional brain connectivity from RPMC to LPMC and LPMC to LPFC, as shown in the Figure 4B. Here, increase in the CoV of the RPMC to LPMC and decrease in the CoV of LPMC to LPFC was related to an increase in the CoV of the FLS score. Also, an effect of the interaction between the skill level and the simulator technology was found on the CoV of the directed functional brain connectivity from LPMC to RPMC as shown in Figure 3C. Here, experts adept at bimanual FLS task performance in physical simulator reduced the CoV of the information flow from LPMC to RPMC in VR simulator compared to physical simulator possibly due to switching to independently controlled bimanual task ^86^ in the unfamiliar VR environment. In contrast, novices were likely performing an independently controlled bimanual task in both the physical and the VR simulators, so an increase in the CoV of the information flow from LPMC to RPMC in the VR simulator may be related to the greater variability of sensory input ^87^ in VR than the physical simulator. Therefore, portable brain imaging provided insights into the effect of interaction between the skill level and the simulator technology related to the interhemispheric inhibition for motor control.

We also found hemispheric lateralization in our right-handed subjects where the coupling between the LPMC and the RPFC (see Figure 2D and 2F) may be related to the detection (efferent copy LPMC to RPFC) and response (cognitive control RPFC to LPMC) to unexpected stimuli ^88^ in the VR environment. Here, higher involvement of RPFC in VR simulator may be related to higher demands on monitoring and checking ^89^. In contrast, the involvement of LPFC (see Figure 2) as the recipient of the corollary discharge information from SMA may be related to its role in analyzing external information during planning goal hierarchy ^89^. Then, any conflict between the efferent information and the sensory reafferent information can lead to a loss of subjective sense of control in the angular gyrus ^13^ (see Figure 1B) that is relevant to evaluate the design of simulator technology. It is postulated that the interaction between the angular gyrus (AG) and the middle frontal gyrus (MFG) is underpinned by the dorsal superior longitudinal fascicle (SLF II) ^90^ where the subjective sense of control may be facilitated by neuroimaging guided transcranial electrical stimulation ^91^ of the AG-MFG interactions ^92^. The dorsal branch of the superior longitudinal fasciculus, responsible for visuospatial integration and motor planning, is found linked to the lateralized hand preference and manual specialization ^93^. Here, the right MFG has been proposed to be a site of convergence of the dorsal and ventral attention networks ^63^ for cognitive control that is relevant in the perception-action cycle. The ventral superior longitudinal fascicle (SLF III) ^90^ is postulated to be more relevant in perception (see Figure 1B) from supramarginal gyrus (SMG) where left MFG and left inferior frontal gyrus (IFG) are more involved in more perceptually demanding FLS tasks, e.g., FLS suturing with intracorporal knot tying ^94^. Here, the ventral stream of perception can be facilitated by neuroimaging guided transcranial electrical stimulation ^91^ of the SMG-IFG interactions ^92^. Then, the coupling between the SMA and LPFC may be related to patterns of pre-learned behavior performed in familiar environments ^88^ in the case of experts in the physical simulator. Here, it is postulated that the interaction between the preSMA/SMA and the PFC/IFG is underpinned by the extended frontal aslant tract (exFAT) ^95^ of the short frontal lobe connections ^96^ that has a role in executive function/ability ^97^. The exFAT may be left lateralized ^95^ that aligns well with left lateralized activation for more complex bimanual FLS tasks, e.g., FLS suturing with intracorporal knot tying ^94^. Although FLS pattern cutting task is also a bimanual task but we only investigated the first sliding window of 54 sec across five repeated trials of FLS tasks when the cutting is with the right hand for all right-handed subjects (cutting direction and sometimes the hand switched at different timepoints after 54 sec due to the surgical field constraints; see FLS pattern cutting video in the supplementary materials). So, we aimed to capture the initial step in FLS pattern cutting skill acquisition to investigate the action-perception link ^55^ when the cognitive-perceptual model ^4^ is developed that is postulated to be lateralized where action=>perception has left hemispheric lateralization while perception=>action has right hemispheric lateralization. We found that the cognitive-action information flow from the right PFC and the efference copy from the left PMC (right handed subjects) was transmitted to the left PFC via SMA as the hub – postulated to underpin action=>perception – that was affected by the skill level (Figure 1C). However, the simulation technology solely affected the cognitive-action information flow from the right PFC to the left PMC while the interaction between the simulation technology and the skill level affected the efference information flow from the left PMC to the right PFC and from the SMA to the left PFC – postulated to capture efference copy and corollary discharge respectively (Figure 1C), which may be related to predictive internal signaling ^62^.

Limitations of this study include the spatial resolution of fNIRS and the optimality of the parameter of the sliding window method for measuring dynamic functional connectivity ^48^. The smallest window greater than 50 sec was found by running stationarity tests on fNIRS time series. Here, a tradeoff was made; on the one hand, the width must be long enough to provide good frequency resolution, while on the other hand, the width must be short enough to satisfy the condition of stationarity. So, instead of an ad-hoc window size ^98^, we searched for an optimal^99^ sliding window pertinent to our data. Also, due to limitations with the spatial resolution of our fNIRS device, we investigated only five brain regions, including LPFC, RPFC, LPMC, RPMC, SMA. Here, the premotor and motor areas were combined in the PMC (see Table S1) and fNIRS optode montage could not distinguish SMA proper from preSMA brain regions that may be important to better assess the temporal structure ^100^ of the perception-action coupling link ^55^. Also, we did not investigate all the subregions of PFC, e.g., ventrolateral PFC and IFG, that may have essential functional interactions during FLS surgical skill acquisition ^91^ where the feasibility of fNIRS’s temporal resolution needs to shown in the future to capture the fast interactions that are expected via shorter frontal lobe connections ^96^.

## Material and Methodology

### 4.1 Subjects and experimental design

The human study was approved by the Institutional Review Board of the Massachusetts General Hospital, University at Buffalo, and the Rensselaer Polytechnic Institute, USA. Seven experienced right-handed surgeons (experts, 5th-year residents, and attending surgeons) and six right-handed medical students (novices, 1st- to 3rd-year residents) participated in the study. The subject details are provided in Table 1. Only right-handed subjects were selected to avoid dominant hemisphere-related inter-subject variability.

**Table 1:**
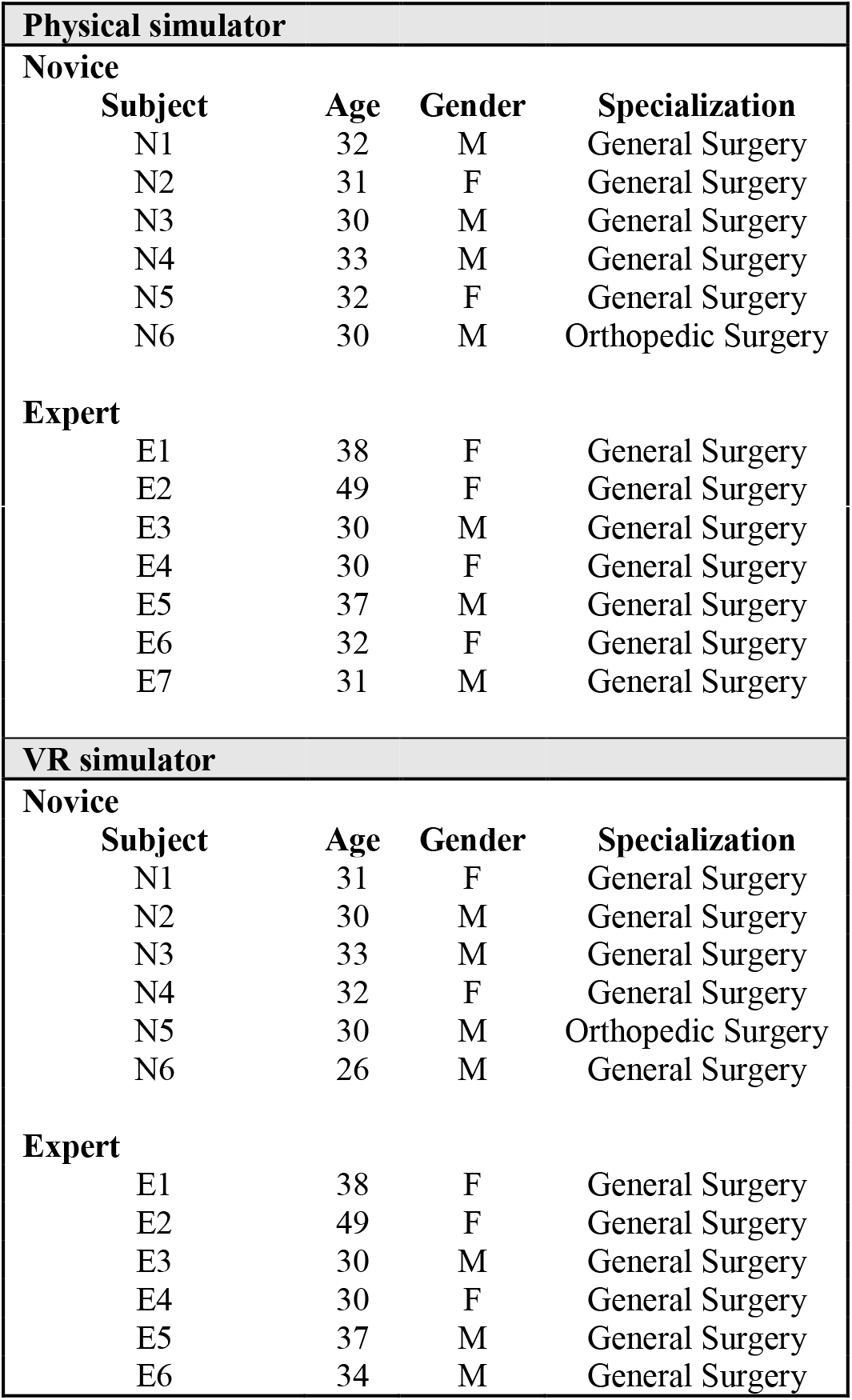
Subject demographics.

| Physical simulator |  |  |  |
| --- | --- | --- | --- |
| Novice |  |  |  |
| Subject | Age | Gender | Specialization |
| N1 | 32 | M | General Surgery |
| N2 | 31 | F | General Surgery |
| N3 | 30 | M | General Surgery |
| N4 | 33 | M | General Surgery |
| N5 | 32 | F | General Surgery |
| N6 | 30 | M | Orthopedic Surgery |
| Expert |  |  |  |
| E1 | 38 | F | General Surgery |
| E2 | 49 | F | General Surgery |
| E3 | 30 | M | General Surgery |
| E4 | 30 | F | General Surgery |
| E5 | 37 | M | General Surgery |
| E6 | 32 | F | General Surgery |
| E7 | 31 | M | General Surgery |
| **VR simulator** |  |  |  |
| **Novice** |  |  |  |
| **Subject** | **Age** | **Gender** | **Specialization** |
| N1 | 31 | F | General Surgery |
| N2 | 30 | M | General Surgery |
| N3 | 33 | M | General Surgery |
| N4 | 32 | F | General Surgery |
| N5 | 30 | M | Orthopedic Surgery |
| N6 | 26 | M | General Surgery |
| **Expert** |  |  |  |
| E1 | 38 | F | General Surgery |
| E2 | 49 | F | General Surgery |
| E3 | 30 | M | General Surgery |
| E4 | 30 | F | General Surgery |
| E5 | 37 | M | General Surgery |
| E6 | 34 | M | General Surgery |

Written consent was obtained from each subject before starting the study. All the subjects were instructed verbally with a standard set of instructions on how to complete the FLS pattern cutting task on the FLS-certified physical and the VR simulator ^101^. For the completion of the FLS pattern cutting task, the right-handed subjects were asked to grasp the gauze using the left grasper (for traction) and cut along (and within) the circular stamp with the right laparoscopic scissors (for cutting). The trial time starts when the subject touches the gauge and ends when the circular cut piece is removed from the gauge frame, and the participants were asked to cut the marked piece of gauze as quickly and as accurately as possible. The data collection was performed with a block design of rest and stimulus period (pattern cutting task) where after a rest period of 1 min (baseline data), the FLS pattern cutting task had to be completed or stopped within 5 min (task data). This was repeated five times (5 trials) for each participant in the repeated-measure study. The performance score for each trial was recorded based on the FLS metrics.

A 32-channel continuous-wave near-infrared spectrometer (CW6 system, TechEn Inc., USA) was used for the optical brain imaging using infrared light at 690nm and 830nm. The optode montage consisted of eight long-distance and eight-short distance sources coupled to 16 detectors. Twenty-five long-distance (30-40mm) channels and eight short-distance (~8mm) channels measured brain activation and systemic physiological signals, respectively (brain regions listed in Table S1 in supplementary materials) that were assessed using the photon migration simulation in the AtlasViewer software ^102^. Here, the photon migration forward matrix represents the sensitivity profile. We selected the average functional near-infrared spectroscopy (fNIRS) signal of the left and the right middle frontal gyrus for the prefrontal cortex activation, i.e., LPFC and RPFC, the left and the right precentral gyrus for the premotor/motor cortex activation, i.e., LPMC and RPMC, and the bilateral supplementary motor area complex for the supplementary motor area activation, i.e., SMA. Table S1 in supplementary materials provides the Montreal Neurological Institute and Hospital (MNI) coordinates. The optical fibers were duly arranged in a cap so that they do not obstruct the free movement of the participant during FLS task performance.

### 4.2 fNIRS data processing for the oxyhemoglobin time-series

Motion artifact detection and correction were performed using Savitzky-Golay filtering ^103^ and band-pass filtering (0.01-0.1Hz) in HOMER3 software (https://github.com/BUNPC/Homer3). Then, modified Beer-Lambert law was used to convert detectors’ optical signals into changes in the oxyhemoglobin (HbO2) concentrations for the partial path-length factors of 6.4 (690nm) and 5.8 (830nm). The short separation channels (inter-optode distance of 8mm) captured the systemic physiological signals originating from non-cortical superficial regions. The averaged signal from the long separation channels (inter-optode distance of 30-40mm) measured the HbO2 changes at each of the following brain regions, LPFC, RPFC, LPMC, RPMC, SMA. Figure S5 in the supplementary materials shows an illustrative plot of the HbO2 time series.

### 4.3 Granger Causality analysis

Granger causality measured the directed functional connectivity that provided the strength and direction of cortical information flow ^54^ between a pair of brain regions from LPFC, RPFC, LPMC, RPMC, SMA ^104^. Granger causality is grounded upon the postulate that one “causally” connected region would leave a component of its signal on another region with some latency, i.e., an autoregressive model (Granger Causality Description provided in the supplementary materials). Here, Short-Time Fourier Transformation (STFT) was employed for nonparametric spectral Granger causality to estimate sliding-window pairwise measures of Granger causality, thereby eliminating the need of explicit autoregressive modeling ^54^. The lowest frequency of 0.02Hz was found from the fNIRS power spectral density (after 0.01-0.1Hz band-pass filtering), so a non-overlapping fixed window size of 54 sec (greater than 50 sec) was selected heuristically ^105^.

### 4.4 Directed functional brain network

Granger causality in the neurovascular frequency range of 0.01 Hz to 0.07Hz ^64^ was used to obtain directed connectivity for each pair of regions (total 20 connections). Figure S1 in the supplementary materials shows an example of 20 inter-regional directed connections for an illustrative time window. The directed connectivity between each pair of brain regions was used to form the directed functional brain network at each time window. Here, selecting the neurovascular frequency range of 0.01 Hz to 0.07Hz ^64^ acts as a filter for systemic and physiological noise like heartbeat and respiration ^106^.

### 4.5 Statistical analysis

The Shapiro-Wilk test was used to test normality for each of the dependent variables (i.e., inter-regional directed functional connectivity metric). Table S8 in the supplementary materials shows the results from Shapiro-Wilk’s test of normality for the Granger causality measure of all each pairs of brain regions in expert and novice while performing FLS task in the physical and virtual simulators across five trials. Then, the directed functional connectivity (Granger causality) between each pair of brain regions for the first window (54 sec) of each trial was used to conduct a repeated-measure two-way multivariate analysis of variance (two-way MANOVA) in SPSS version 27 (IBM, USA) to determine whether there is a significant difference in the inter-regional directed functional connectivity based on the skill level (expert, novice), simulator technology (physical simulator, VR simulator) and their interaction. We conducted two-way MANOVA in SPSS version 27 (IBM, USA) to determine whether there is a significant difference in the coefficient of variation (CoV) of inter-regional directed functional connectivity across trials based on the skill level, simulator technology, and their interaction. We conducted a repeated-measure two-way analysis of variance (ANOVA) in SPSS version 27 (IBM, USA) to determine whether there is a significant difference in the FLS score based on the skill level, simulator technology, and their interaction. We also conducted a repeated-measure two-way analysis of variance (ANOVA) in SPSS version 27 (IBM, USA) to determine whether there is a significant difference in the CoV of FLS score based on the skill level, simulator technology, and their interaction. The Levene test was used to test the homogeneity of variance. All the significance levels were set at alpha=0.05. To determine how the dependent variables (i.e., inter-regional directed functional connectivity) differ for the independent variables – the skill level, simulator technology, and their interaction, i.e., the tests of between-subjects effects, alpha with multiple comparison correction (False Discovery Rate) and partial eta squared effect size were used. Then, we conducted brain-behavior analysis via multiple regression (backward elimination with a probability of F for removal ≥ 0.1) in SPSS version 27 (IBM, USA) to find the relationship of the inter-regional directed functional brain connectivity with the FLS score. Then, in SPSS Amos (IBM, USA), the path analysis was performed from the the skill level (expert, novice) and simulator technology (physical simulator, VR simulator) to the dependent variables, inter-regional directed functional brain connectivity leading to the dependent variable, FLS score.

## Acknowledgment

The authors gratefully acknowledge the support of this work through the Medical Technology Enterprise Consortium (MTEC) award #W81XWH2090019 (2020-628), and the U.S. Army Futures Command, Combat Capabilities Development Command Soldier Center STTC cooperative research agreement #W912CG-21-2-0001.

## Disclosures

Authors have neither relevant financial or competing interests nor other potential conflicts of interest.

## Author contributions

Conceptualization, A.D.; methodology, A.K., X.I., S.D., A.D.; software, A.K.; validation, A.K., X.I., S.D., A.D.; formal analysis, A.K., A.D.; investigation, A.K., A.D.; resources, S.D.; data curation, A.K.; writing—original draft preparation, A.K., A.D.; writing—review and editing, B.M., J.N., S.S., X.I., S.D., A.D.; visualization, A.K., A.D.; supervision, X.I., S.D., A.D.; project administration, S.D.; funding acquisition, B.M., J.N., S.S., X.I., S.D., A.D. All authors have read and agreed to the published version of the manuscript.

## Notes

### Competing Interest Statement

The authors have declared no competing interest.

## Reference

1. Riener, R. & Harders, M. Virtual Reality in Medicine. (Springer-Verlag, 2012). doi:10.1007/978-1-4471-4011-5.

2. Fuster, J. M. Chapter 8 - Prefrontal Cortex in Decision-Making: The Perception-Action Cycle. in Decision Neuroscience (eds. Dreher, J.-C. & Tremblay, L.) 95–105 (Academic Press, 2017). doi:10.1016/B978-0-12-805308-9.00008-7.

3. Renner, R. S., Velichkovsky, B. M. & Helmert, J. R. The perception of egocentric distances in virtual environments - A review. ACM Comput. Surv. 46, 23:1–23:40 (2013).

4. Cioffi, D. Beyond attentional strategies: cognitive-perceptual model of somatic interpretation. Psychol Bull 109, 25–41 (1991).

5. Marucci, M. et al. The impact of multisensory integration and perceptual load in virtual reality settings on performance, workload and presence. Sci Rep 11, 4831 (2021).

6. Wolpert, D. M. & Miall, R. C. Forward Models for Physiological Motor Control. Neural Netw 9, 1265–1279 (1996).

7. Kawato, M. Internal models for motor control and trajectory planning. Curr Opin Neurobiol 9, 718–727 (1999).

8. Sommer, M. A. & Wurtz, R. H. Brain circuits for the internal monitoring of movements. Annu Rev Neurosci 31, 317–338 (2008).

9. Christensen, A. et al. An intact action-perception coupling depends on the integrity of the cerebellum. J Neurosci 34, 6707–6716 (2014).

10. Hurley, S. Perception and Action: Alternative Views. Synthese 129, 3–40 (2001).

11. Levac, D. E., Huber, M. E. & Sternad, D. Learning and transfer of complex motor skills in virtual reality: a perspective review. Journal of NeuroEngineering and Rehabilitation 16, 121 (2019).

12. Haar, S., Donchin, O. & Dinstein, I. Individual Movement Variability Magnitudes Are Explained by Cortical Neural Variability. J. Neurosci. 37, 9076–9085 (2017).

13. Haggard, P. Sense of agency in the human brain. Nat Rev Neurosci 18, 196–207 (2017).

14. Poldrack, R. A. et al. The Neural Correlates of Motor Skill Automaticity. J Neurosci 25, 5356–5364 (2005).

15. Yücel, M. A., Selb, J. J., Huppert, T. J., Franceschini, M. A. & Boas, D. A. Functional Near Infrared Spectroscopy: Enabling Routine Functional Brain Imaging. Curr Opin Biomed Eng 4, 78–86 (2017).

16. Nemani, A. et al. Assessing bimanual motor skills with optical neuroimaging. Science Advances 4, eaat3807 (2018).

17. Birkmeyer, J. D. et al. Surgical Skill and Complication Rates after Bariatric Surgery. New England Journal of Medicine 369, 1434–1442 (2013).

18. Dehabadi, M., Fernando, B. & Berlingieri, P. The use of simulation in the acquisition of laparoscopic suturing skills. International Journal of Surgery 12, 258–268 (2014).

19. Voorhorst, F., Meijer, D., Overbeeke, C. & Smets, G. Depth perception in laparoscopy through perception-action coupling. Minimally Invasive Therapy & Allied Technologies 7, 325–334 (1998).

20. Bahrami, P. et al. Functional MRI-compatible laparoscopic surgery training simulator. Magnetic Resonance in Medicine 65, 873–881 (2011).

21. Roberts, K. E., Bell, R. L. & Duffy, A. J. Evolution of surgical skills training. World Journal of Gastroenterology 12, 3219–3224 (2006).

22. Kunert, W. et al. Learning curves, potential and speed in training of laparoscopic skills: a randomised comparative study in a box trainer. Surg Endosc 35, 3303–3312 (2021).

23. Wright, W. G. Using virtual reality to augment perception, enhance sensorimotor adaptation, and change our minds. Front. Syst. Neurosci. 8, (2014).

24. Sigrist, R., Rauter, G., Riener, R. & Wolf, P. Augmented visual, auditory, haptic, and multimodal feedback in motor learning: a review. Psychon Bull Rev 20, 21–53 (2013).

25. Ritter, E. M. & Scott, D. J. Design of a proficiency-based skills training curriculum for the fundamentals of laparoscopic surgery. Surg Innov 14, 107–112 (2007).

26. Grantcharov, T. P. & Funch-Jensen, P. Can everyone achieve proficiency with the laparoscopic technique? Learning curve patterns in technical skills acquisition. Am J Surg 197, 447–449 (2009).

27. Wolpert, D. M., Miall, R. C. & Kawato, M. Internal models in the cerebellum. Trends Cogn. Sci. (Regul. Ed.) 2, 338–347 (1998).

28. Ebner, T. J. Cerebellum and Internal Models. in Handbook of the Cerebellum and Cerebellar Disorders 1279–1295 (Springer, Dordrecht, 2013). doi:10.1007/978-94-007-1333-8_56.

29. Badre, D. & D’Esposito, M. Is the rostro-caudal axis of the frontal lobe hierarchical? Nature Reviews Neuroscience vol. 10 659–669 (2009).

30. Badre, D. & D’Esposito, M. Is the rostro-caudal axis of the frontal lobe hierarchical? Nature Reviews Neuroscience 10, 659–669 (2009).

31. Badre, D. Cognitive control, hierarchy, and the rostro–caudal organization of the frontal lobes. Trends in Cognitive Sciences 12, 193–200 (2008).

32. Koechlin, E. & Summerfield, C. An information theoretical approach to prefrontal executive function. Trends Cogn Sci 11, 229–235 (2007).

33. Christoff, K. & Gabrieli, J. D. E. The frontopolar cortex and human cognition: Evidence for a rostrocaudal hierarchical organization within the human prefrontal cortex. Psychobiology 28, 168–186 (2000).

34. Milner, A. D. How do the two visual streams interact with each other? Exp Brain Res 235, 1297–1308 (2017).

35. Tanji, J., Okano, K. & Sato, K. C. Neuronal activity in cortical motor areas related to ipsilateral, contralateral, and bilateral digit movements of the monkey. J Neurophysiol 60, 325–343 (1988).

36. Debaere, F., Wenderoth, N., Sunaert, S., Van Hecke, P. & Swinnen, S. P. Cerebellar and premotor function in bimanual coordination: parametric neural responses to spatiotemporal complexity and cycling frequency. Neuroimage 21, 1416–1427 (2004).

37. Swinnen, S. P. & Wenderoth, N. Two hands, one brain: cognitive neuroscience of bimanual skill. Trends Cogn Sci 8, 18–25 (2004).

38. Ohuchida, K. et al. The frontal cortex is activated during learning of endoscopic procedures. Surgical Endoscopy (2009) doi:10.1007/s00464-008-0316-z.

39. Leff, D. R., Orihuela-Espina, F., Leong, J., Darzi, A. & Yang, G.-Z. Modelling dynamic fronto-parietal behaviour during minimally invasive surgery--a Markovian trip distribution approach. Med Image Comput Comput Assist Interv 11, 595–602 (2008).

40. Wanzel, K. R. et al. Visual-spatial ability and fMRI cortical activation in surgery residents. Am J Surg 193, 507–510 (2007).

41. Leff, D. R., Orihuela-Espina, F., Atallah, L., Darzi, A. & Yang, G.-Z. Functional near infrared spectroscopy in novice and expert surgeons--a manifold embedding approach. Med Image Comput Comput Assist Interv 10, 270–277 (2007).

42. Gao, Y. et al. Decreasing the Surgical Errors by Neurostimulation of Primary Motor Cortex and the Associated Brain Activation via Neuroimaging. Front Neurosci 15, 651192 (2021).

43. Leff, D. R. et al. Functional prefrontal reorganization accompanies learning-associated refinements in surgery: a manifold embedding approach. Comput Aided Surg 13, 325–339 (2008).

44. Khoe, H. C. H. et al. Use of prefrontal cortex activity as a measure of learning curve in surgical novices: results of a single blind randomised controlled trial. Surg Endosc 34, 5604–5615 (2020).

45. Gao, Y. et al. Functional Brain Imaging Reliably Predicts Bimanual Motor Skill Performance in a Standardized Surgical Task. IEEE Trans Biomed Eng 68, 2058–2066 (2021).

46. Kaminski, M. J. & Blinowska, K. J. A new method of the description of the information flow in the brain structures. Biol. Cybern. 65, 203–210 (1991).

47. Kamiński, M., Ding, M., Truccolo, W. A. & Bressler, S. L. Evaluating causal relations in neural systems: granger causality, directed transfer function and statistical assessment of significance. Biol Cybern 85, 145–157 (2001).

48. Hutchison, R. M. et al. Dynamic functional connectivity: Promise, issues, and interpretations. Neuroimage 80, 10.1016/j.neuroimage.2013.05.079 (2013).

49. Kao, C.-H. et al. Functional brain network reconfiguration during learning in a dynamic environment. Nat Commun 11, 1682 (2020).

50. Seth, A. K., Barrett, A. B. & Barnett, L. Granger causality analysis in neuroscience and neuroimaging. Journal of Neuroscience 35, 3293–3297 (2015).

51. Heitger, M. H. et al. Motor learning-induced changes in functional brain connectivity as revealed by means of graph-theoretical network analysis. NeuroImage 61, 633–650 (2012).

52. Gerraty, R. T., Davidow, J. Y., Wimmer, G. E., Kahn, I. & Shohamy, D. Transfer of learning relates to intrinsic connectivity between hippocampus, ventromedial prefrontal cortex, and large-scale networks. J Neurosci 34, 11297–11303 (2014).

53. Strangman, G. E., Li, Z. & Zhang, Q. Depth Sensitivity and Source-Detector Separations for Near Infrared Spectroscopy Based on the Colin27 Brain Template. PLoS One 8, e66319 (2013).

54. Dhamala, M., Rangarajan, G. & Ding, M. Analyzing information flow in brain networks with nonparametric Granger causality. NeuroImage 41, 354–362 (2008).

55. Latash, M. L. Efference copy in kinesthetic perception: a copy of what is it? Journal of Neurophysiology 125, 1079–1094 (2021).

56. Raos, V. & Savaki, H. E. The Role of the Prefrontal Cortex in Action Perception. Cerebral Cortex 27, 4677–4690 (2017).

57. Fuster, J. M. The Prefrontal Cortex—An Update: Time Is of the Essence. Neuron 30, 319–333 (2001).

58. Lebedev, M. A., Douglass, D. K., Moody, S. L. & Wise, S. P. Prefrontal cortex neurons reflecting reports of a visual illusion. J Neurophysiol 85, 1395–1411 (2001).

59. Rolls, E. T., Huang, C.-C., Lin, C.-P., Feng, J. & Joliot, M. Automated anatomical labelling atlas 3. NeuroImage 206, 116189 (2020).

60. Cona, G. & Semenza, C. Supplementary motor area as key structure for domain-general sequence processing: A unified account. Neurosci Biobehav Rev 72, 28–42 (2017).

61. Dutta, A. et al. Interhemispheric functional connectivity in the primary motor cortex distinguishes between training on a physical and a virtual surgical simulator. bioRxiv 2021.07.10.451831 (2021) doi:10.1101/2021.07.10.451831.

62. Straka, H., Simmers, J. & Chagnaud, B. P. A New Perspective on Predictive Motor Signaling. Curr Biol 28, R232–R243 (2018).

63. Japee, S., Holiday, K., Satyshur, M. D., Mukai, I. & Ungerleider, L. G. A role of right middle frontal gyrus in reorienting of attention: a case study. Frontiers in Systems Neuroscience 9, 23 (2015).

64. Nemani, A. et al. Functional brain connectivity related to surgical skill dexterity in physical and virtual simulation environments. Neurophotonics 8, (2021).

65. Sadato, N., Yonekura, Y., Waki, A., Yamada, H. & Ishii, Y. Role of the Supplementary Motor Area and the Right Premotor Cortex in the Coordination of Bimanual Finger Movements. J Neurosci 17, 9667–9674 (1997).

66. Toyokura, M., Muro, I., Komiya, T. & Obara, M. Relation of bimanual coordination to activation in the sensorimotor cortex and supplementary motor area: analysis using functional magnetic resonance imaging. Brain Res Bull 48, 211–217 (1999).

67. Ullén, F., Forssberg, H. & Ehrsson, H. H. Neural networks for the coordination of the hands in time. J Neurophysiol 89, 1126–1135 (2003).

68. Gerloff, C., Corwell, B., Chen, R., Hallett, M. & Cohen, L. G. Stimulation over the human supplementary motor area interferes with the organization of future elements in complex motor sequences. Brain 120 (Pt 9), 1587–1602 (1997).

69. Mushiake, H., Fujii, N. & Tanji, J. Visually guided saccade versus eye-hand reach: contrasting neuronal activity in the cortical supplementary and frontal eye fields. J Neurophysiol 75, 2187–2191 (1996).

70. Pierrot-Deseilligny, C., Israël, I., Berthoz, A., Rivaud, S. & Gaymard, B. Role of the different frontal lobe areas in the control of the horizontal component of memory-guided saccades in man. Exp Brain Res 95, 166–171 (1993).

71. Steyvers, M. et al. High-frequency transcranial magnetic stimulation of the supplementary motor area reduces bimanual coupling during anti-phase but not in-phase movements. Exp Brain Res 151, 309–317 (2003).

72. Euston, D. R., Gruber, A. J. & McNaughton, B. L. The Role of Medial Prefrontal Cortex in Memory and Decision Making. Neuron 76, 1057–1070 (2012).

73. Fuster, J. M. Upper processing stages of the perception–action cycle. Trends in Cognitive Sciences 8, 143–145 (2004).

74. Prasad, R., Muniyandi, M., Manoharan, G. & Chandramohan, Servarayan. M. Face and Construct Validity of a Novel Virtual Reality-Based Bimanual Laparoscopic Force-Skills Trainer With Haptics Feedback. Surg Innov 25, 499–514 (2018).

75. Koechlin, E., Ody, C. & Kouneiher, F. The architecture of cognitive control in the human prefrontal cortex. Science 302, 1181–1185 (2003).

76. Jenkins, I. H., Brooks, D. J., Nixon, P. D., Frackowiak, R. S. J. & Passingham, R. E. Motor sequence learning: A study with positron emission tomography. Journal of Neuroscience 14, 3775–3790 (1994).

77. Bates, J. F. & Goldman-Rakic, P. S. Prefrontal connections of medial motor areas in the rhesus monkey. J Comp Neurol 336, 211–228 (1993).

78. Morecraft, R. J. & Van Hoesen, G. W. Frontal granular cortex input to the cingulate (M3), supplementary (M2) and primary (M1) motor cortices in the rhesus monkey. J Comp Neurol 337, 669–689 (1993).

79. Makoshi, Z., Kroliczak, G. & van Donkelaar, P. Human supplementary motor area contribution to predictive motor planning. J Mot Behav 43, 303–309 (2011).

80. McCloskey, D. I. Corollary Discharges: Motor Commands and Perception. in Comprehensive Physiology 1415–1447 (American Cancer Society, 2011). doi:10.1002/cphy.cp010232.

81. Abe, M. et al. Functional Coupling of Human Prefrontal and Premotor Areas during Cognitive Manipulation. J. Neurosci. 27, 3429–3438 (2007).

82. Willingham, D. B. A Neuropsychological Theory of Motor Skill Learning. Psychological Review 105, 558–584 (1998).

83. Thorp, E. B., Kording, K. P. & Mussa-Ivaldi, F. A. Using noise to shape motor learning. Journal of Neurophysiology 117, 728–737 (2017).

84. Wu, H. G., Miyamoto, Y. R., Castro, L. N. G., Ölveczky, B. P. & Smith, M. A. Temporal structure of motor variability is dynamically regulated and predicts motor learning ability. Nat Neurosci 17, 312–321 (2014).

85. Ranganathan, R. & Newell, K. M. Emergent flexibility in motor learning. Exp Brain Res 202, 755–764 (2010).

86. Fling, B. W. & Seidler, R. D. Task-Dependent Effects of Interhemispheric Inhibition on Motor Control. Behav Brain Res 226, 211–217 (2012).

87. Trommershäuser, J., Gepshtein, S., Maloney, L. T., Landy, M. S. & Banks, M. S. Optimal compensation for changes in task-relevant movement variability. J Neurosci 25, 7169–7178 (2005).

88. MacNeilage, P. F., Rogers, L. J. & Vallortigara, G. ORIGINS OF THE Left & Right Brain. Scientific American 301, 60–67 (2009).

89. Kaller, C. P., Rahm, B., Spreer, J., Weiller, C. & Unterrainer, J. M. Dissociable Contributions of Left and Right Dorsolateral Prefrontal Cortex in Planning. Cerebral Cortex 21, 307–317 (2011).

90. Nakajima, R., Kinoshita, M., Shinohara, H. & Nakada, M. The superior longitudinal fascicle: reconsidering the fronto-parietal neural network based on anatomy and function. Brain Imaging and Behavior 14, 2817–2830 (2020).

91. Walia, P., Kumar, K. N. & Dutta, A. Neuroimaging Guided Transcranial Electrical Stimulation in Enhancing Surgical Skill Acquisition. Comment on Hung et al. The Efficacy of Transcranial Direct Current Stimulation in Enhancing Surgical Skill Acquisition: A Preliminary Meta-Analysis of Randomized Controlled Trials. Brain Sci. 2021, 11, 707. Brain Sciences 11, 1078 (2021).

92. Walia, P., Kamat, A., De, S. & Dutta, A. Dynamic causal modeling for EEG during complex laparoscopic skill acquisition. (2021).

93. Howells, H. et al. Frontoparietal Tracts Linked to Lateralized Hand Preference and Manual Specialization. Cerebral Cortex (New York, NY) 28, 2482 (2018).

94. Walia, P. et al. Neuroimaging guided tES to facilitate complex laparoscopic surgical tasks – insights from functional near-infrared spectroscopy. (2021) doi:10.21203/rs.3.rs-730076/v1.

95. Pascual-Diaz, S., Varriano, F., Pineda, J. & Prats-Galino, A. Structural characterization of the Extended Frontal Aslant Tract trajectory: A ML-validated laterality study in 3T and 7T. NeuroImage 222, 117260 (2020).

96. Catani, M. et al. Short frontal lobe connections of the human brain. Cortex 48, 273–291 (2012).

97. La Corte, E. et al. The Frontal Aslant Tract: A Systematic Review for Neurosurgical Applications. Frontiers in Neurology 12, 51 (2021).

98. Li, Z. et al. Dynamic functional connectivity revealed by resting-state functional near-infrared spectroscopy. Biomedical Optics Express 6, 2337 (2015).

99. Wang, Z. et al. Best window width determination and glioma analysis application of dynamic brain network measure on resting-state functional magnetic resonance imaging. Journal of Medical Imaging and Health Informatics 6, 1735–1740 (2016).

100. Schwartze, M., Rothermich, K. & Kotz, S. A. Functional dissociation of pre-SMA and SMA-proper in temporal processing. NeuroImage 60, 290–298 (2012).

101. Linsk, A. M. et al. Validation of the VBLaST pattern cutting task: a learning curve study. Surg Endosc 32, 1990–2002 (2018).

102. Aasted, C. M. et al. Anatomical guidance for functional near-infrared spectroscopy: AtlasViewer tutorial. Neurophotonics 2, (2015).

103. Jahani, S., Setarehdan, S. K., Boas, D. A. & Yücel, M. A. Motion artifact detection and correction in functional near-infrared spectroscopy: a new hybrid method based on spline interpolation method and Savitzky–Golay filtering. Neurophotonics 5, (2018).

104. Seth, A. K., Barrett, A. B. & Barnett, L. Granger Causality Analysis in Neuroscience and Neuroimaging. Journal of Neuroscience 35, 3293–3297 (2015).

105. Leonardi, N. & Van De Ville, D. On spurious and real fluctuations of dynamic functional connectivity during rest. Neuroimage 104, 430–436 (2015).

106. Auer, D. P. Spontaneous low-frequency blood oxygenation level-dependent fluctuations and functional connectivity analysis of the ‘resting’ brain. Magnetic Resonance Imaging 26, 1055–1064 (2008).

